# Behavior Variability of a Conditional Gene Knockout Mouse as a Measure of Subtle Phenotypic Trait Expression. The Case of Mouse Executive Function Distortion

**DOI:** 10.1101/229856

**Authors:** Pavel Prosselkov, Qi Zhang, Hiromichi Goto, Denis Polygalov, Thomas J. McHugh, Shigeyoshi Itohara

## Abstract

Task learning relies on brain executive function (EF), the construct of controlling and coordinating behavior under the everlasting flow of environmental changes. We have previously shown, that a complete knockout of a vertebrate brain-specific pair of gene paralogs (*Ntng1/2*) distorts the mouse EF, making behavior less predictable (more variable) via the affected working memory and attention (1). In the current study, conditionally targeting either serotonin transporter (*5-HTT*) or *Emx1*-expressing neurons, we show that the cell type-specific ablation of *Ntng1* within the excitatory circuits of either cortex or thalamus does not have a profound impact on the EF but rather affects its certain modalities, *i.e.* impulsivity and/or selective attention, modulated by cognitive demand. Several mice of both conditional genotypes simultaneously occupy either top or bottom parameter-specific behavioral ranks, indicative of a subject-unique antagonistic either proficit or deficit of function within the same behavior. Employing genotype-attributable behavior variability as a phenotypic trait, we deduce, that *Ntng1*-parsed excitatory pathways contribute but do not fully reconstitute the attention-impulsivity phenotypes, associated with the mouse EF deficit. However, complete knockdown of *Ntng1/2*, and associated with it behavior variability, explains the deficit of executive function and task learning.

## INTRODUCTION

Executive function (EF) can be conceptualised as a measure of general cognitive processes ontogenetic canalization (2). In mice EF is frequently assessed by the 5-choice serial reaction time task (5-CSRTT), a behavioral test of sustained attention parameters (3): 1. response accuracy (success of correct over all attempted trials), interpreted as a measure of spatially divided attention; 2. omission errors (trials with no response made, a correlate of selective attention); 3. premature responses (responses preceding trial onset, as a measure of impulsivity), and 4. erroneous (perseverative) nose pokings (after the outcome feedback) are measures of inhibitory control required for cognitive flexibility, representing approximate constructs of impulsivity and compulsivity, respectively. Additionally, response and reward collection latencies relate to processing speed and/or motivational factors (4). EF strongly depends on cortical and subcortical brain areas interactions primary guided by the thalamus-located somatosensory inputs, *i.e.* top-down and bottom-up information colliding flows, modulated by the cognitive demand (1). Besides, behavioral inhibition (3) and cognitive flexibility (5) are specifically dependent on serotonin presence within thalamus nuclei. Despite traditionally being viewed as a passive relay of sensory information (6; see also 7 for ref.), thalamus integrates information across cortical networks (8), and, via persistent reverberating activity (9-11), bidirectionaly links with all cortical areas (12-15), justifying its role in the EF signals processing as the cortex itself.

Behavioral individuality is an inevitably “unpredictable outcome” of life matter development elaborated on a top of genes and/or environment interactions (16), representing inherited genetic variability (17), and meaning that genetic content is capable to explain (at least partially) an individual’s behavioral (in)variability. At the cellular level, neuronal correlated variability for spikes and subjects’ performance displays the same relationship as learning and attention (18), perhaps through the affected impulsivity (as shown in this study).

Previously, we have shown that a pair of brain vertebrate-specific presynaptically expressed gene paralogs, *NTNG1/2*, is functionally characterised by antagonistic pleiothropy (19), and associated with the human IQ scores (20). Both genes are complementary expressed across the cognition-loaded brain areas with *Ntng1* being predominantly located within bottom-up (sensory) circuits (*e.g.* thalamus and shallower cortical layers) while *Ntng2* - within deeper cortical and subcortical areas (21), referred as top-down circuits. Cognition-deprived phenotypes are produced upon either of *Ntng* paralogs global knockdown with affected working memory and attention (1).

In the current study, to assess the degree of a genetic manipulation impact on the EF behavior perturbations, *Ntng1* was locally ablated in the excitatory cortical (*Emx1*-promoter) and serotonin transporter expressing (*5-HTT*-promoter) thalamic neurons. We demonstrate that behavioral variability, reported as a behavioral rank variance, and viewed as a reporter trait for a specific gene manipulation, can be used to explain the behavior of mouse genetically-mixed population without any other detectable (substantially significant) phenotypic measures exerted by conditional deletion of a gene in a specific brain area, untractable by traditional arithmetic mean-based methods.

## RESULTS

Brain areas spatially-restricted (conditional) ablation of *Ntng1* within excitatory cortical and serotonin transporter expressing thalamic neurons was achieved by the use of *Emx1-cre* and *5-HTT-cre* expressing reporter lines, respectively, as described previously (22). Briefly, upon a recombination with the *Ntng1^ƒ/ƒ^* allele the resulted *Emx1-Ntng1^−/−^* mice displayed reduced in the cortex and no expression of *Ntng1* in the *stratum lacunosum* and outer molecular layer of the hippocampus and piriform cortex layer I comparing to the non-recombined control line. Another line, *5-HTT-Ntng1^−/−^* mice, showed the selective reduction of *Ntng1* in the ventral basal and lateral geniculate nuclei of thalamus and cortical layer IV, when compared with the control (22). Attention and impulsivity (AI) estimate during the 5-CSRTT by comparison of averaged behavior either across the spatials or summed spatials as low and high cognitive demand (LCD and HCD, respectively) sessions (Fig.1) did not reveal any dramatic behavioral changes upon the local gene knockout. Some statistical significance has been reached for the *Emx1-Ntng1^−/−^* during the LCD and *5-HTT-Ntng1^−/−^*mice during the HCD sessions for the percent of success (Sc, Fig.1B-1,2). *5-HTT-Ntng1^−/−^* mice were also better than controls at premature pokings (PreP) but only during the HCD (Fig.1C-2). And both conditional lines were slightly better at erroneous pokings (EP): *Emx1-Ntng1^−/−^* (Fig.1E-1) throughout the all sessions, and *5-HTT-Ntng1^−/−^* during the HCD only (Fig.1E-2). It should be noted, that, in most cases, the knockout mice outperformed their genetically-unmodified littermates (but carrying the *Ntng1^ƒ/ƒ^* allele), including the task learning (spatial 1) for *Emx1-Ntng1^−/−^* subjects (Supplementary Table 2-1 (ST2-1)).

**Figure 1.**
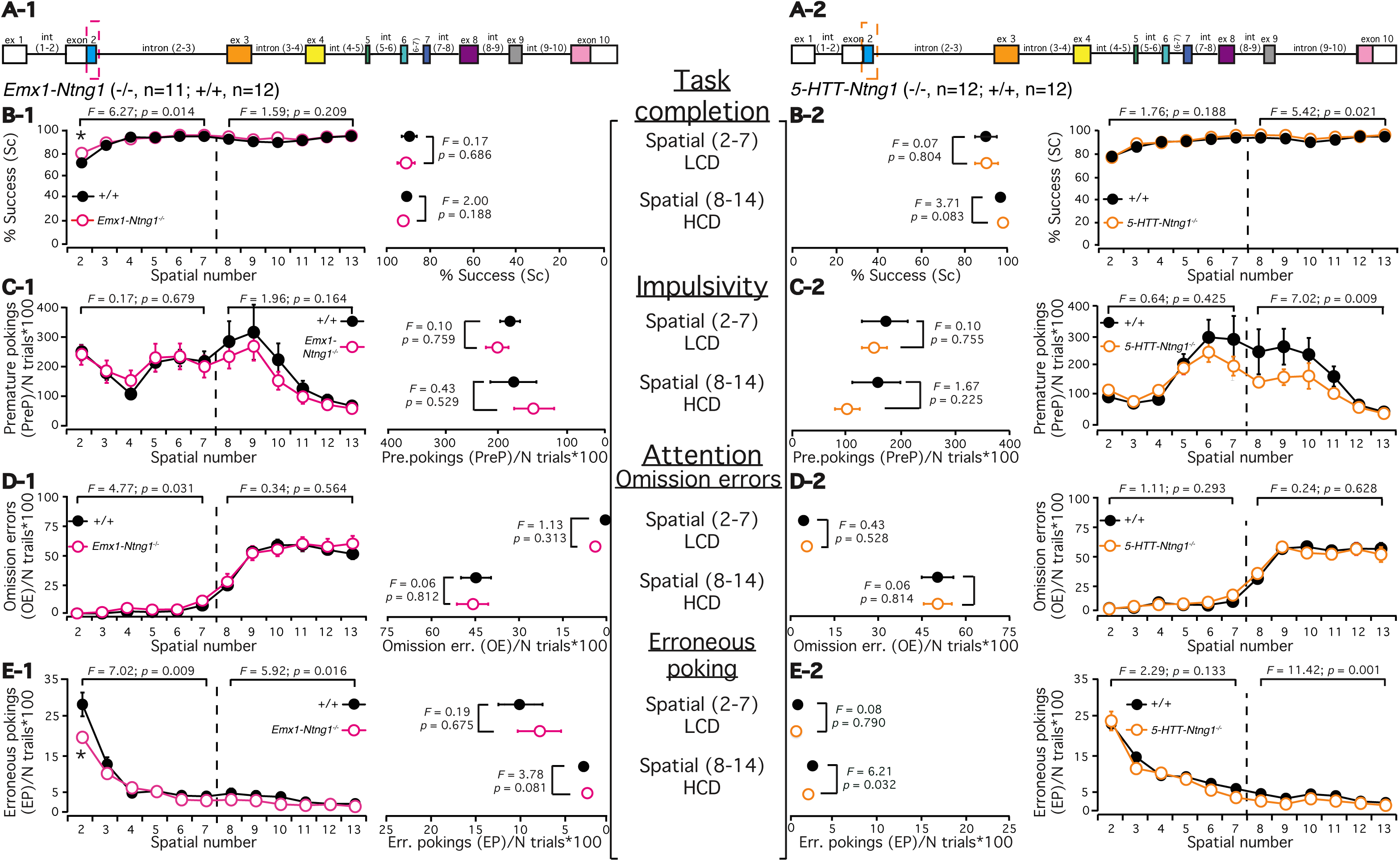
Attention and Impulsivity (AI) estimate for *Emx1-Ntng1^−/−^* and c*Ntng1-5-HTT*^−/−^ by comparison of averaged behavior for 5-CSRTT. **A.** Exon-intron composition of mouse *Ntng1* with the removed part dash-outlined, see (22) for the details of each construct design. (**B-E, outmost left and right**) Mice performance over the spatials (2-13) presented by four behavioral parameters. The dashed line separating spatials (2-7) and (8-13) indicates the data split on the low cognitive demand (LCD) and high cognitive demand (HCD) sessions. (**B-E, middle left and right**) The same data as above but averaged per LCD and HCD sessions. The data for each parameter are presented as a mean±SEM and provided in **ST2-1** (for *Emx1-Ntng1^−/−^*) and **ST2-2** (for *5-HTT-Ntng1^−/−^*). Two-way and one-way ANOVA were used for statistics with *indicating *p*<0.05.

To deduce potential genotype-phenotype inferences we have randomized the behavioral output regardless of the input genotype, as originally described in (1), using rank and calculating its variance, as a proportion of variance explained (PVE), (Fig.2; ST2-1 and ST2-2). No substantial differences neither across genotypes nor LCD/HCD sessions have been observed for the generated ranks but for the genotype-attributable rank variances (Fig.2A,E). *Emx1-Ntng1^−/−^* mice persistently show 3 times lower (Fig.2A-1, E-1) and *5-HTT-Ntng1^−/−^* mice (for HCD only) 2 times higher (Fig.2E-2) rank PVE comparing to the wild type littermates. This indicates that cortical excitatory and thalamic relay neurons differentially affect mouse behavioral variability modulated by the cognitive demand. At the same time, the majority of both knockout mice display a trend of occupying either top or bottom ranks for a single parameter during the LCD (Fig.2B-1: n=5/5, top/bottom for *Emx1-Ntng1^−/−^*; Fig.2B-2: n=4/6, top/bottom for *5-HTT-Ntng1^−/−^*), abolished when the demand gets higher (Fig.2F-1: n=4/1, top/bottom for *Emx-Ntng1^−/−^*; Fig.2F-2: n=0/2, top/bottom for *5HTT-Ntng1^−/−^*). Several mice have been noted to occupy parameter-specific either top or bottom ranking simultaneously, uniquely segregating their abilities within the functional proficit and deficit of the task-specific cognitive abilities (Fig.2C), strongly affected by the HCD (Fig.2G). *Emx1-Ntng1^−/−^*mice showed lower correlations of the prepulse inhibition (PreP) and omission errors (OE) with other parameters regardless of the cognitive pressure (Fig.2D-1, H-1), with *5-HTT-Ntng1^−/−^* mice showing similar feature but only during the HCD (Fig.2D-2, H-2). Thus, several aspects of the AI-related behaviors are shared among the excitatory cortical and thalamic brain areas expressing *Ntng1.*

**Figure 2.**
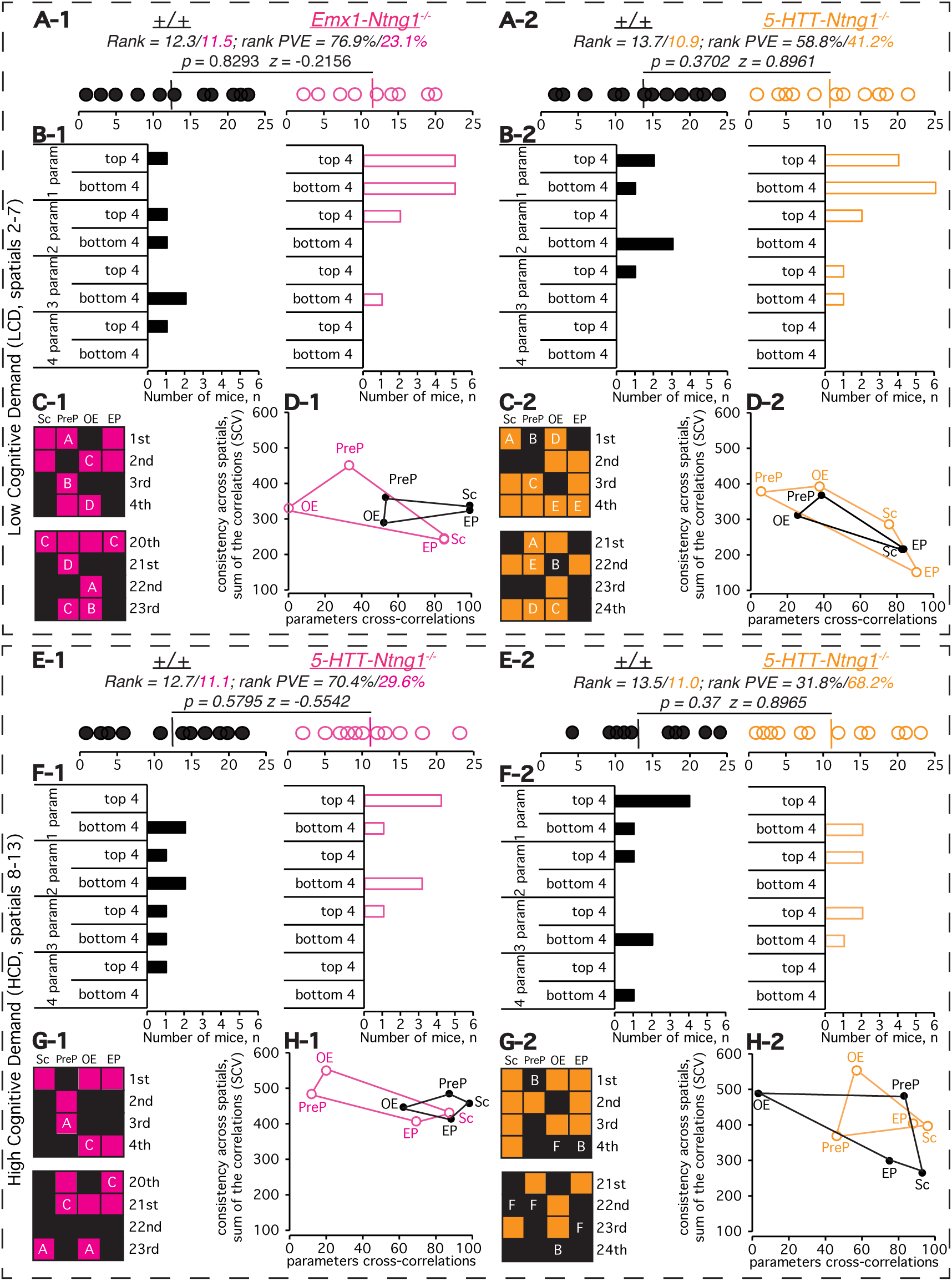
Attention and Impulsivity (AI) estimate by the analysis of rank and its variance for *Emx1-Ntng1^−/−^* and *5-HTT-Ntng1^−/−^* mice performing 5-CSRTT. **A,E.** Mouse ranks and rank PVE (proportion of variance explained) based on four parameters rank measures (**Fig.1**) with all calculations detailed in **ST2-1** (for *Emx1-Ntng1^−/−^*) and **ST2-2** (for *5-HTT-Ntng1^−/−^*). The rank sorting was done in a genotype-independent manner treating all mice equally regardless of the genotype. Ranking for each out of four parameters was done independently of other parameters with a final re-ranking of the ranks sum to generate the final rank (shown). In case of an equal sum of the ranks, the mice were given identical ranks. PVE was calculated as a square of within genotype rank variance divided on the sum of each genotype variances squares multiplied on 100%. **B,F**. Mouse rank distribution across one-to-four parameters as top 4 and bottom 4 performers. **C,G.** Genotype-specific placing among the mice. An alphabetic character inside a square (from A to F) denotes a mouse simultaneously occupying a top four and bottom four ranks. **D,H.** Behavioral consistency of mice across the sessions (*y* axis, sum of *r^2^* correlations of a single session ranks *vs.* final ranks for each mouse across the sessions) and behavioral parameter cross-correlations (*x* axis, the *r^2^* correlation of a parameter final ranking *vs.* final ranking for all 4 parameters). The gene ablation-specific phenotype severity can be assessed visually by matching each parameter-corresponding verteces of the obtained quadruples. *p* value was obtained by Wilcoxon rank sum test.

Behavioral phenotypic proximity between the genotypes supporting genotype-phenotype relationships have been assessed in two ways. First, by the *C*-means fuzzy clustering using the phenotypic data input as a rank to predict the corresponding genotypes as an output (Fig.3A), and second, as a geometrical function of the liner distance between the genotypes when a parameter-specific rank is plotted against its PVE (Fig.3B). Clustering probability of the genotype-specific segregation is the strongest during HCD for *Emx1-Ntng1^−/−^* mice (Fig.3A-1, lower panel), and during LCD for *5HTT-Ntng1^−/−^* population (Fig.3A-2, higher panel), indicative of the potential role switches between the somatosensory and cortical/subcortical excitatory pathways when the cognitive demand to focus is augmented. In overall, the low resolution for the genotypes clustering probabilities, obtained by the first method, resembles that of the experimental data (Fig.2A,E), and is further supported by the second method. It indicates that the phenotypic distance within the *5-HTT-Ntng1^−/−^* population is reduced during the HCD (Fig.3B-2), but stays nearly the same for the *Emx1-Ntng1^−/−^* mice (Fig.3B-1), probably due to the most affected parameter switch (PreP to OE in the latter case).

**Figure 3.**
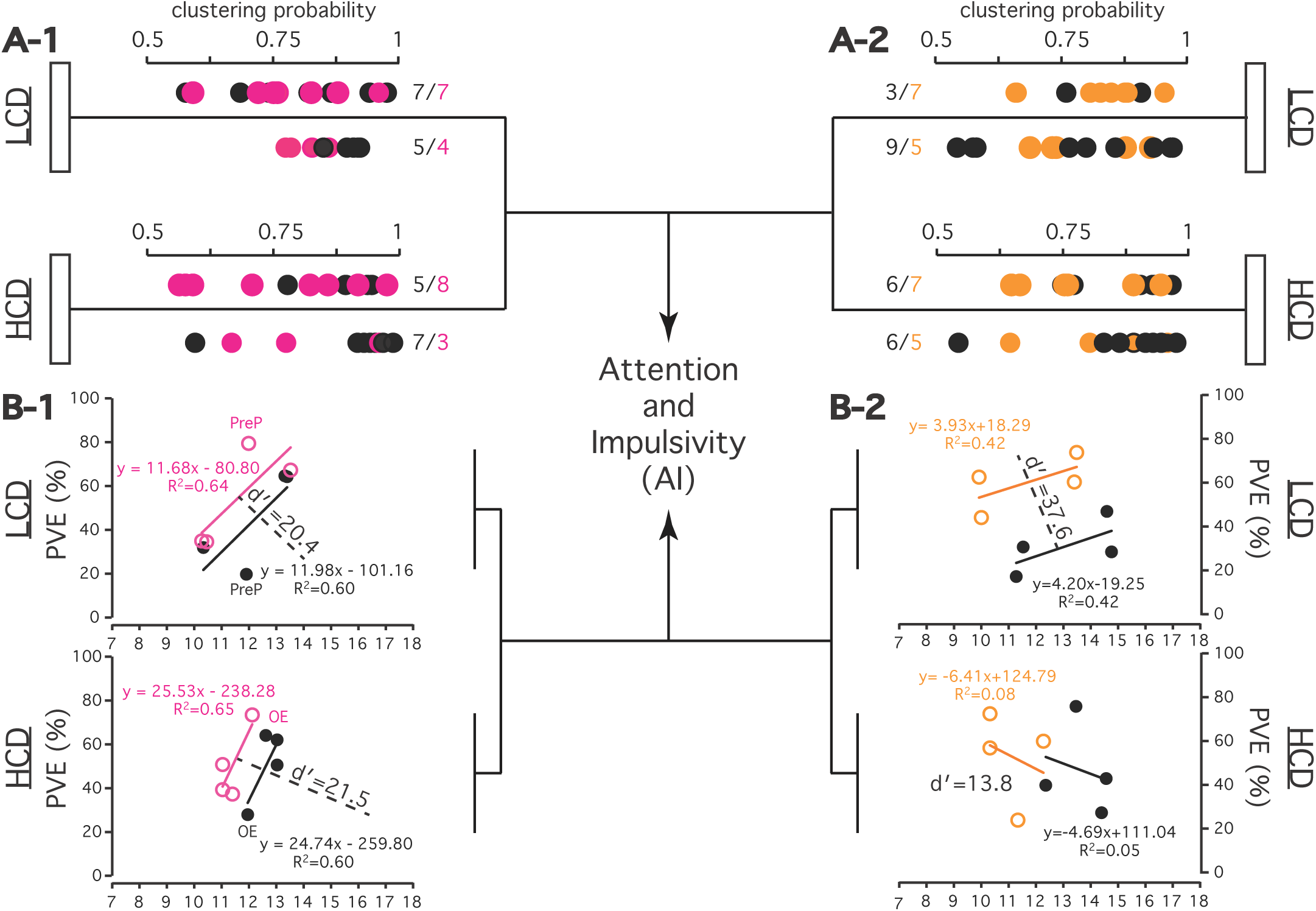
Behavioral phenotypic proximity assessment for *Emx1-Ntng1^−/−^* and *5-HTT-Ntng1^−/−^* genotypes and their wild type littermates by two means. genotype predictions by *C*-means fuzzy clustering (**A**), and by linear regression plot of genotype-specific rank PVE *vs.* rank (**B**). **A**. Causal genotype-phenotype relationship deduction. *C*-means fuzzy clustering (Euclidean *C*-means) was done in a genotype-blind manner using the obtained behavioral ranks (Fig.2) to predict subjects’ genotype, as described in (1). See ST3 for the exact values of clustering probabilities. **B.** Descriptive proximity of the attention and impulsivity (AI) phenotypes for the *Emx1-Ntng1^−/−^* and *5-HTT-Ntng1^−/−^* mice and their wild type littermates. Phenotypic distance between the genotypes (*d’*, dashed line) is presented geometrically by the linear equation, calculated as *d’=|c_2_-c_1_|*, where *ax+by+c=0* (Euclidean geometry). Rank (*x* coordinates) and PVE values (*y* coordinates) are from Fig.2A,E, ST2-1 and ST2-2.

To extend the search for a correlate explaining the obtained behavioral pattern (Fig.2A,E), we have probed another measure, capable to absorb both independently-descriptive genotype-specific variables, such as parameter-specific rank and its PVE (ST2-1 and ST2-2), by multiplying them (Fig.4A). This allowed us to build patterns, expressed as bars, comparing each gene contribution (as a knockout) against other gene content, and to compare different genes of distinct functions (as knockouts) tested under similar conditions (*e.g.* LCD or HCD). As it can be seen from Fig.4A, PreP during LCD, and OE during both, LCD and HCD, are two parameters, the most affected by *Ntng1* ablation within *Emx1* and *5-HTT*-expressing neurons. Similar approach has been used to re-analyze the already reported datasets (1; ST4-1 and ST4-2), incorporating data for the globally ablated *Ntng1/2* paralogs (Fig.4B,C), and complementing the current conditional knockout and already published findings (1). Further integrating all generated parameters (ST2-1, and ST2-2) into the global ranks (Fig.5A), we conclude that serotonin plays a substantial role in the AI deficit during HCD and operant conditioning learning throughout the whole process. Adding to that, working memory deficit associated with the *Ntng1/2* ablation (1), supports the observed operant conditioning learning difficulties by the *Ntng1/2* global knockout mice (Fig.5B, ST4-1, and ST4-2), and complements the AI deficit (Fig.5C, ST5-1, and ST5-2).

**Figure 4.**
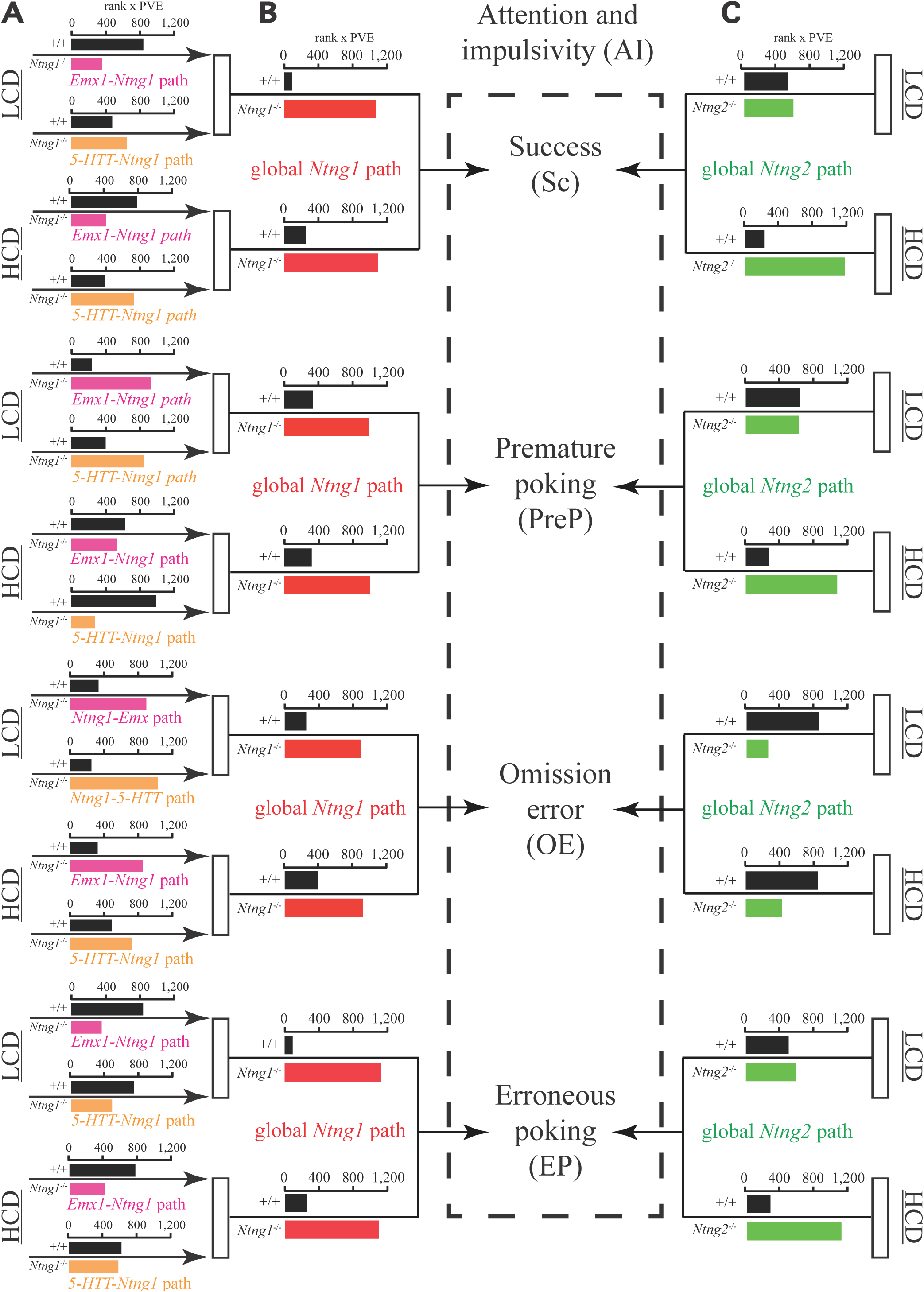
Effect of partial (A, pink/orange) or global (B,C red/green) *Ntng* paralogs ablation on the AI parameters. (Sc, PreP, OE, and EP) as measured by the 5-CSRTT, presented against the total gene content (+/+) and calculated as a parameter-specific rank x PVE (Fig.2); **B** and **C** data are from (1): Fig.1,2). For the calculus details see ST2-1, ST2-2, ST4-1 and ST4-2.

**Figure 5.**
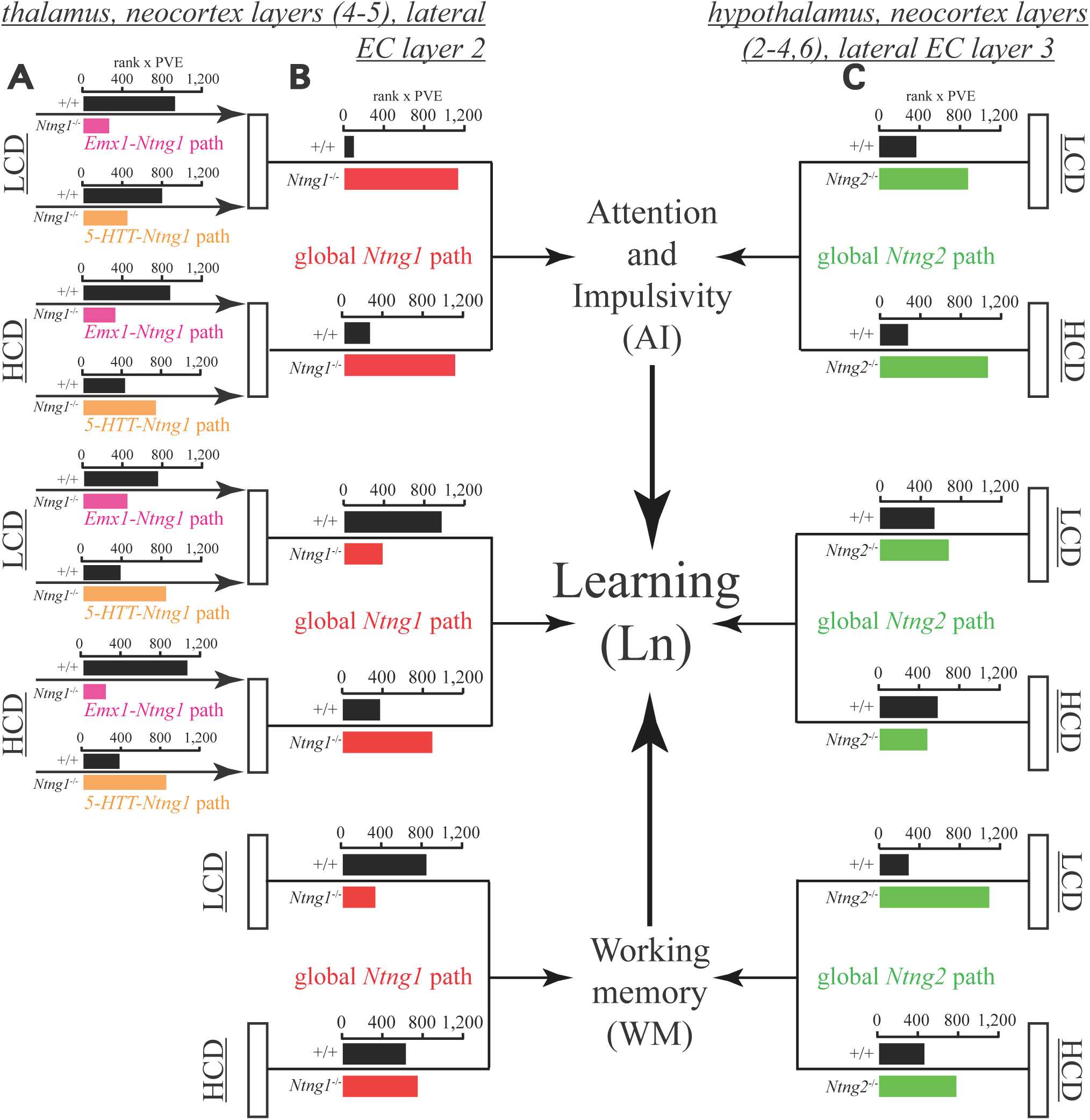
Operant conditioning learning (Ln) association with AI and WM parsed by *Ntng* gene paralogs’ brain expression pathways, interrogated either by global (B,C) or partial ablation (A). The “rank x PVE” represent global rank sum values based on the parameters-specific rank and PVE values from Fig.4. For the calculus details see Fig.2, ST2-1, ST2-2, ST4-1, ST4-2, ST5-1, and ST5-2. The top panel (in *italics*) indicates the brain areas complementary expressing *Ntng* gene paralogs (1).

## DISCUSSION

To assess the degree of *Ntng1* gene local ablation on the EF of mouse brain, we have prepared two mouse lines with the expression of *Ntng1* spatially turned off in excitatory cortical and subcortical *Emx1* and serotonin transporter-expressing thalamic relay neurons. No behavioral data have revealed any major impact on the mouse behavior for the 5-CSRTT (Fig.1). However, employing the previously used approach (1), substantial genotype-phenotype inferences were obtained by ranking the randomized behavioral output with the mouse input genotype kept blind. Parameter-specific deficits have been deduced such as affected OE and PreP (Figs.2-3), as effective measures of AI (3), respectively. The elevated impulsivity observed for the knockout mice is frequently reported as a core endophenotypic trait for many psychiatric disorders characterised by impulsive action and choices dependent on serotonin (23), exacerbated by extended inter-trial intervals (delay periods) that are challenging but necessary for proper motor preparations (9,11), and instrumental for the correct responses (10). The reported associations of the human gene ortholog *NTNG1* with schizophrenia (19-20) underscores this gene function as filtering or sensory information gating (it is strongly expressed in thalamus and cortex, 22), and explaining the observed mouse phenotypes by a disregulation of a flow of sensory information affecting the cognitive functions.

As anticipated, the degree of the obtained phenotype is mild comparing to the complete *Ntng* paralogs caused behaviors (1) and does not fully explain it (Fig.5). One potential reason for this would be the omitted by this project importance of inhibitory neurons, with the graded control of thalamic output by the inhibitory systems known to match ongoing behavioral demands (14). *Ntng1* is expressed in inhibitory hippocampal GABAergic neurons, and in amygdala it is mainly detected in the lateral nucleus and *zona incerta* (22). Careful assessment of the inhibitory neurons role in the EF control would be a tantalizing target for future projects.

Based on the rhetoric assumption that directionality of a phenotypic trait (its advantage or disadvantage, to acquire deficit or proficit, *i.e.* trait attenuation or augmentation) is only descriptive, strictly context-dependent and does not explicitly express the underlying gene phenotypic function, we proposed that phenotypic variability would be sufficiently able to explain genotype-phenotype interactions of a genetically heterogeneous population. This is opposite to the classical approach when two groups of distinct genotypes (*i.e.* knockouts and their genetically-unmodified siblings) are matched against each other behavioral scores in search for a significantly different task-specific parameter for a gene function-related conclusion claimed to be causal enough to explain the observed behavioral deficit.

Behavioral variability within a population of mice can be either affected by a gene targeted knockout or not, eventually being translated into the genotype-specific rank PVE, attributable to the specifically induced genotype. In a search for the obtained results explanation we have chosen an approach emphasizing the importance of behavioral variability as a trait affected by the gene manipulation. The given method is strongly biased (gives advantage) towards the cases characterised by the larger variance and lower ranks (as a poor performance) but complements the commonly used approach of the means comparison where behavioral variability among the mice is historically considered as the worst factor of all. And it is especially valuable in the absence of obviously evident phenotypes reaching a “statistical significance” to be reported as per current scientific publication standards (24).

The generated graphs (rank multiplied on PVE, Figs.4-5) represent a kind of a behavioral pattern for a population of genetically heterogeneous mice for the specific task. A similar behavioral pattern but within a distinct behavioral context (*e.g.* another behavioral paradigm) or another animal population underwent identical genetic perturbations would indicate a stable genotype-phenotype correlative interactions. The obtained pattern does not explain the ablated gene function in terms of phenotypic advantages-disadvantages but rather characterises a distance from the other gene content presented by the non-modified (wild type) littermates. The presented method allows for cross-population and cross-paradigm mouse behavioral assessments comparing it with the other gene content provided by the wild type littermates or another line of genetically modified subjects.

Focusing on the variability itself as a behavioral trait, it is interesting to note, that in human subjects neural variability represents an individual trait exhibiting distinct behavioral capabilities (25), explainable by a variance, and that “intelligence test scores estimate a relative rank order rather than measure the construct of intelligence” (26). Since we have shown that behavioral variability is able (at least partially) to explain the deficit in the EF (Figs.4-5), and it is known that the variance in EF control is attributable only to cognitive performance variance, *i.e.* general intelligence (27), behavioral variability is able to fulfill the function of a cognitive ability-describing trait.

## CONCLUSION

A local area gene-ablation caused behavioral variability can be used to deduce the genotype-phenotype specific interactions affecting the executive function of the brain. Assessing the degree of a genotype-attributable behavioral variance engagement into a cognitive task allows to make an inference regarding to a subtle phenotypic trait expression affected by a local area gene manipulation. Behavioral variability of a genetically heterogeneous population of mice can be used as an estimate of the comprising it homogenous subpopulation cognitive abilities.

## MATERIALS AND METHODS

### Animals handling, behavioral set-up, and knockout lines generation

Animal rearing and handling were performed in accordance with the guidelines of RIKEN Institutional Animal Care and Experimentation Committee (ethics approval number H29-2-235(3)). The behavioral set-ups are as described in (1), and the knockout lines were generated as per (22). In brief, *Ntng1^ƒ/ƒ^* mice were crossed with the *Emx1-Cre* line followed by triggered recombination in excitatory neurons of cerebral cortex and limbic structures. Upon crossing with the *5-HTT-Cre*, the recombination took place in excitatory neurons of the first-order thalamic relay and raphe nuclei. *Emx1-Ntng1^−/−^* and *5-HTT-Ntng1^−/^* mouse pups were born and developed normally with no detectable anatomical or in-cage behavioral abnormalities when matched to the *Ntng1^ƒ/ƒ^* littermates. In *Emx1-Ntng1^−/−^* mouse brains *Ntng1* expression was dramatically decreased in the cortex but stayed unaltered in the thalamus, with the immune signal also lowered in the layer 1a of piriform cortex, the terminals for the mitral cell axons of the olfactory buls. In *5-HTT-Ntng1^−/−^* mice *Ntng1* expression was unaltered in the cortex but significantly decreased in the ventral basal, lateral geniculate, and medial geniculate nuclei of the thalamus with a marked decrease in the encoded netrin-G1 protein presence in layer 4 of the cortex.

### Data analysis, definition of LCD and HCD, fuzzy *C*-Means Clustering

All raw data including the generated ranks and the proportion of variance explained (PVE) calculations, cell-embedded formulas and graphs are presented in ST2-ST5. *Ntng1/2* global knockout datasets (ST4-ST5) are from (1) with the “rank x PVE” values used to compare with the conditional knockout mice bevaior. While performing the 5-CSRTT the attentional load was incremented by a shorter cue duration and longer inter-trial intervals, later denoted as a low cognitive demand (LCD, spatials 2-7), and higher cognitive demand (HCD, spatials 8-13). Fuzzy *C*-Means clustering represents a reverse task of predicting a subjects’ cluster (genotype in our case) based on the input data (behavioral ranks) under the genotype-blind input conditions. Further details can be found in (1) Supplementary materials.

### Statistics

One and two-way ANOVA was calculated using StatPlus (AnalystSoft Inc.). Wilcoxon rank sum test was done by Matlab (v.7.9.0 2009b) by the function *ranksum.*

## SUPPLEMENTARY MATERIALS

Contain Supplementary Tables (ST2-ST5), as Excel files except ST3 provided as a pdf file.

## ACKNOWLEDGEMENTS

This work was in part supported by the “Funding Program for World-Leading Innovative R&D on Science and Technology (FIRST Program)” initiated by the Council for Science and Technology Policy (CSTP), and KAKENHI 15H04290 from the Japan Society for the Promotion of Science (JSPS).

## COMPETING INTERESTS

Authors would like to express a lack of any competing interests associated with the work.

---

**Supplementary Table 3 (ST3).** *C*-means fuzzy clustering (Euclidean *C*-means) rank probabilities for *Emx1*-*Ntng1^−/−^* and *5-HTT*-*Ntng1^−/−^* mice performance on the 5-choice serial reaction time task (5-CSRTT).

| MouseID | X1 | X2 | Cem.cluster |
| --- | --- | --- | --- |
| 1292 | 0.971194328 | 0.028805672 | 1 |
| 1236 | 0.954745496 | 0.045254504 | 1 |
| 1222 | 0.935816244 | 0.064183756 | 1 |
| 1235 | 0.870727133 | 0.129272867 | 1 |
| 1301 | 0.857281879 | 0.142718121 | 1 |
| 1228 | 0.815973987 | 0.184026013 | 1 |
| 1300 | 0.810241352 | 0.189758648 | 1 |
| 1248 | 0.746420223 | 0.253579777 | 1 |
| 1234 | 0.738442275 | 0.261557725 | 1 |
| 1256 | 0.72935276 | 0.27064724 | 1 |
| 1223 | 0.706827886 | 0.293172114 | 1 |
| 1303 | 0.669912252 | 0.330087748 | 1 |
| 1272 | 0.572814641 | 0.427185359 | 1 |
| 1225 | 0.558424095 | 0.441575905 | 1 |
| 1302 | 0.083735977 | 0.916264023 | 2 |
| 1243 | 0.087782277 | 0.912217723 | 2 |
| 1271 | 0.097168259 | 0.902831741 | 2 |
| 1259 | 0.110884603 | 0.889115397 | 2 |
| 1296 | 0.148071512 | 0.851928488 | 2 |
| 1257 | 0.159000963 | 0.840999037 | 2 |
| 1280 | 0.182422526 | 0.817577474 | 2 |
| 1281 | 0.225845049 | 0.774154951 | 2 |
| 1254 | 0.236862407 | 0.763137593 | 2 |

| QC Measure | Cem_QC |
| --- | --- |
| fhv | 0.201770755 |
| apd | 9.315988268 |
| pd | 9.545188185 |
| xb | 0.010251666 |
| fs | -25.7575624 |
| pc | 0.715021771 |
| pe | 0.44828755 |
| pre | 95.85991388 |
| si | 0.24185849 |

| MouseID | X1 | X2 | Cem.cluster |
| --- | --- | --- | --- |
| 1910 | 0.967003554 | 0.032996446 | 1 |
| 1846 | 0.944971779 | 0.055028221 | 1 |
| 1912 | 0.934603138 | 0.065396862 | 1 |
| 1838 | 0.928213393 | 0.071786607 | 1 |
| 1891 | 0.902810891 | 0.097189109 | 1 |
| 1908 | 0.894528843 | 0.105471157 | 1 |
| 1907 | 0.888669072 | 0.111330928 | 1 |
| 1857 | 0.765215407 | 0.234784593 | 1 |
| 1868 | 0.753286749 | 0.246713251 | 1 |
| 1890 | 0.746830575 | 0.253169425 | 1 |
| 1833 | 0.742325733 | 0.257674267 | 1 |
| 1861 | 0.655569113 | 0.344430887 | 1 |
| 1870 | 0.637823967 | 0.362176033 | 1 |
| 1897 | 0.023219431 | 0.976780569 | 2 |
| 1837 | 0.041497705 | 0.958502295 | 2 |
| 1895 | 0.054164734 | 0.945835266 | 2 |
| 1869 | 0.071068015 | 0.928931985 | 2 |
| 1855 | 0.086636718 | 0.913363282 | 2 |
| 1834 | 0.110940888 | 0.889059112 | 2 |
| 1875 | 0.139456452 | 0.860543548 | 2 |
| 1853 | 0.172120404 | 0.827879596 | 2 |
| 1911 | 0.201383692 | 0.798616308 | 2 |
| 1859 | 0.364905462 | 0.635094538 | 2 |
| 1888 | 0.471137599 | 0.528862401 | 2 |

| QC Measure | Cem_QC |
| --- | --- |
| fhv | 0.179795196 |
| apd | 10.81906165 |
| pd | 10.70943992 |
| xb | 0.007041227 |
| fs | -35.9943064 |
| pc | 0.753577808 |
| pe | 0.396166088 |
| pre | 117.7803381 |
| si | 0.214449133 |

| MouseID | X1 | X2 | Cem.cluster |
| --- | --- | --- | --- |
| 1292 | 0.952557717 | 0.047442283 | 1 |
| 1248 | 0.95250862 | 0.04749138 | 1 |
| 1302 | 0.921191833 | 0.078808167 | 1 |
| 1301 | 0.912376999 | 0.087623001 | 1 |
| 1235 | 0.893132879 | 0.106867121 | 1 |
| 1300 | 0.867606202 | 0.132393798 | 1 |
| 1228 | 0.831201648 | 0.168798352 | 1 |
| 1236 | 0.794162541 | 0.205837459 | 1 |
| 1303 | 0.748313679 | 0.251686321 | 1 |
| 1234 | 0.675428503 | 0.324571497 | 1 |
| 1254 | 0.553431251 | 0.446568749 | 1 |
| 1281 | 0.538143087 | 0.461856913 | 1 |
| 1223 | 0.52546059 | 0.47453941 | 1 |
| 1222 | 0.034828418 | 0.965171582 | 2 |
| 1271 | 0.055404196 | 0.944595804 | 2 |
| 1296 | 0.063382807 | 0.936617193 | 2 |
| 1243 | 0.071471007 | 0.928528993 | 2 |
| 1256 | 0.082116167 | 0.917883833 | 2 |
| 1257 | 0.094728317 | 0.905271683 | 2 |
| 1259 | 0.10800378 | 0.89199622 | 2 |
| 1280 | 0.254796411 | 0.745203589 | 2 |
| 1272 | 0.366776706 | 0.633223294 | 2 |
| 1225 | 0.441907542 | 0.558092458 | 2 |

| QC Measure | Cem_QC |
| --- | --- |
| fhv | 0.214193892 |
| apd | 12.47660286 |
| pd | 12.44663823 |
| xb | 0.008197161 |
| fs | -31.28506848 |
| pc | 0.734994733 |
| pe | 0.418329232 |
| pre | 104.4172345 |
| si | 0.248889369 |

| MouseID | X1 | X2 | Cem.cluster |
| --- | --- | --- | --- |
| 1875 | 0.945717956 | 0.054282044 | 1 |
| 1891 | 0.941602693 | 0.058397307 | 1 |
| 1908 | 0.907049915 | 0.092950085 | 1 |
| 1868 | 0.899748813 | 0.100251187 | 1 |
| 1837 | 0.848866171 | 0.151133829 | 1 |
| 1869 | 0.828158797 | 0.171841203 | 1 |
| 1910 | 0.768078402 | 0.231921598 | 1 |
| 1833 | 0.733215453 | 0.266784547 | 1 |
| 1912 | 0.706012221 | 0.293987779 | 1 |
| 1907 | 0.698158601 | 0.301841399 | 1 |
| 1834 | 0.652429984 | 0.347570016 | 1 |
| 1838 | 0.542100557 | 0.457899443 | 1 |
| 1888 | 0.535252651 | 0.464747349 | 1 |
| 1853 | 0.502343617 | 0.497656383 | 1 |
| 1890 | 0.070435428 | 0.929564572 | 2 |
| 1895 | 0.118797886 | 0.881202114 | 2 |
| 1870 | 0.146632357 | 0.853367643 | 2 |
| 1859 | 0.152009202 | 0.847990798 | 2 |
| 1861 | 0.179949674 | 0.820050326 | 2 |
| 1846 | 0.201998198 | 0.798001802 | 2 |
| 1897 | 0.223035615 | 0.776964385 | 2 |
| 1857 | 0.272382844 | 0.727617156 | 2 |
| 1911 | 0.374268744 | 0.625731256 | 2 |
| 1855 | 0.376556769 | 0.623443231 | 2 |

| QC Measure | Cem_QC |
| --- | --- |
| fhv | 0.426415625 |
| apd | 7.296242585 |
| pd | 6.605961542 |
| xb | 0.012399117 |
| fs | -21.12556049 |
| pc | 0.675782558 |
| pe | 0.495315436 |
| pre | 84.47706536 |
| si | 0.2510265 |

